# Force-independent interactions of talin and vinculin govern integrin-mediated mechanotransduction

**DOI:** 10.1101/629683

**Authors:** Paul Atherton, Franziska Lausecker, Alexandre Carisey, Andrew Gilmore, David Critchley, Igor Barsukov, Christoph Ballestrem

## Abstract

Talin, vinculin and paxillin are core components of the dynamic link between integrins and actomyosin. Here we study the mechanisms that mediate their activation and association using a mitochondrial-targeting assay, structure-based mutants, and advanced microscopy. As expected, full-length vinculin and talin are auto-inhibited and do not interact with each other in this state. Contrary to previous models that propose a critical role for forces driving talin-vinculin association, our data show that force-independent relief of auto-inhibition is sufficient to mediate their tight interaction. Interestingly, paxillin can bind to both talin and vinculin when either is inactive. Further experiments demonstrate that adhesions containing paxillin and vinculin can form without talin following integrin activation. However, these are largely deficient in exerting traction forces to the matrix. Our observations lead to a model whereby paxillin contributes to talin and vinculin recruitment into nascent adhesions. Activation of the talin-vinculin axis subsequently leads to the engagement with the traction force-machinery and focal adhesion maturation.

## Introduction

Focal adhesions (FAs) are sites of integrin-mediated cell adhesion to the extracellular matrix (ECM). The abundance and diversity of proteins in FAs (Horton et al., 2015) allows FAs to act as efficient signaling hubs, regulating multiple aspects of cell behavior including migration, differentiation and proliferation (Geiger and Yamada, 2011). Talin and vinculin are two critical regulators of the mechanical link between integrins and the actin cytoskeleton (Gauthier and Roca-Cusachs, 2018). Structurally, both talin (Goult et al., 2013a) and vinculin (Chorev et al., 2018; Cohen et al., 2005) exist in closed (auto-inhibited) and open conformations. This has led to an attractive model in which actomyosin-mediated forces are envisaged to induce conformational changes that unmask binding sites in both proteins that support their mutual interaction and association with the contractile actomyosin machinery, plus other binding partners (Chorev et al., 2018; del Rio et al., 2009; Sun et al., 2017; Yao et al., 2014; Yao et al., 2016). For vinculin, force is thought to overcome the strong auto-inhibitory interaction (Kd 0.1 µM (Cohen et al., 2005)) between the globular N-terminal head (domains D1-D4) and the C-terminal D5 tail domain (Vt) that masks the talin binding site in the D1 domain (Cohen et al., 2005). Furthermore, a FRET conformation sensor has shown that vinculin is in an open conformation within FAs (Chen et al., 2005). For talin, the primary auto-inhibitory interaction is between the F3 domain of the N-terminal FERM domain and R9, one of the 13 α-helical bundles (R1-R13) in the flexible C-terminal rod (Calderwood et al., 2013). Although a role for force in relieving talin auto-inhibition is less clear than for vinculin, in vitro studies have clearly shown that force acting on individual talin rod domains can unmask vinculin binding sites (VBS) buried within their cores (del Rio et al., 2009; Yao et al., 2014), thereby facilitating vinculin binding (Carisey et al., 2013; Yao et al., 2016). FRET-based tension sensors for both talin and vinculin show that both are under tension within FAs (Austen et al., 2015; Grashoff et al., 2010; Kumar et al., 2016; LaCroix et al., 2018), and myosin-dependent stretching of talin has been demonstrated in cells (Margadant et al., 2011). Together, these experiments suggest a model where actomyosin-mediated forces activate talin and promote vinculin binding, strengthening engagement of talin with the actomyosin machinery which is critical for the transmission of force from the cytoskeleton to the ECM via FAs (Atherton et al., 2016; Atherton et al., 2015; Elosegui-Artola et al., 2016; Goult et al., 2018; Sun et al., 2017). However, the idea that force induces activation of both proteins largely derives from in vitro biochemical experiments, and a clear understanding of the processes involved in talin and vinculin activation in cells requires further investigation. The vinculin/talin axis forms a scaffold for many adhesion proteins during FA development, including the signaling protein paxillin (Carisey et al., 2013). Conversely, paxillin, which can bind to both vinculin (Deakin et al., 2012) and talin (Zacharchenko et al., 2016), is also implicated in recruiting vinculin to adhesions downstream of myosin-dependent tyrosine phosphorylation (Pasapera et al., 2010). However, to what extent paxillin binding to talin or vinculin is dependent on their activation states remains unclear.

In this study, we aimed to determine the contribution of force to the talin/vinculin interaction by targeting proteins to the mitochondria, reducing the complexity of the environment. We combined this approach with structure-based talin and vinculin point and deletion mutants to reveal the contribution of specific talin domains towards talin activation and subsequent vinculin binding and show that disrupting auto-inhibition of either molecule is sufficient to induce a very stable force-independent interaction. Interestingly, the adhesion protein paxillin can be recruited to both talin and vinculin in their inactive forms independently of force, leading to a model where force-independent processes initiate an adhesion complex including integrin, talin, vinculin and paxillin that subsequently engages the actomyosin machinery resulting in reinforcement of this linkage.

## Results

The enormous complexity of protein-protein associations within FAs makes it virtually impossible to analyse molecular rearrangements and to separate force-dependent and force-independent processes. To overcome these limitations, and to test the role of forces in vinculin and talin activation, we fused the C-terminus of talin or vinculin with the mitochondrial targeting sequence from the C-terminus of BAK (cBAK: aa1072-1162) (Fig. 1a). Co-localisation with the mitochondrial-specific dye Mitotracker established successful targeting of both constructs to the mitochondria (Fig. 1b). Neither integrins nor filamentous actin were found next to mitochondria (Fig 1b and 1c), confirming a force-free environment (Fig 1b and 1c, (Detmer and Chan, 2007)). Crucially, neither cBAK-fused wild-type vinculin (vinFL-cBAK), nor wild-type talin (talinFL-cBAK) recruited either co-expressed or endogenous talin or vinculin respectively (Fig. 1d). This suggests that the C-terminal cBAK tag does not affect the structure, function or activation status of wild-type talin or vinculin, and that an essential signal required for the talinvinculin association is absent from mitochondria.

**Fig. 1.**
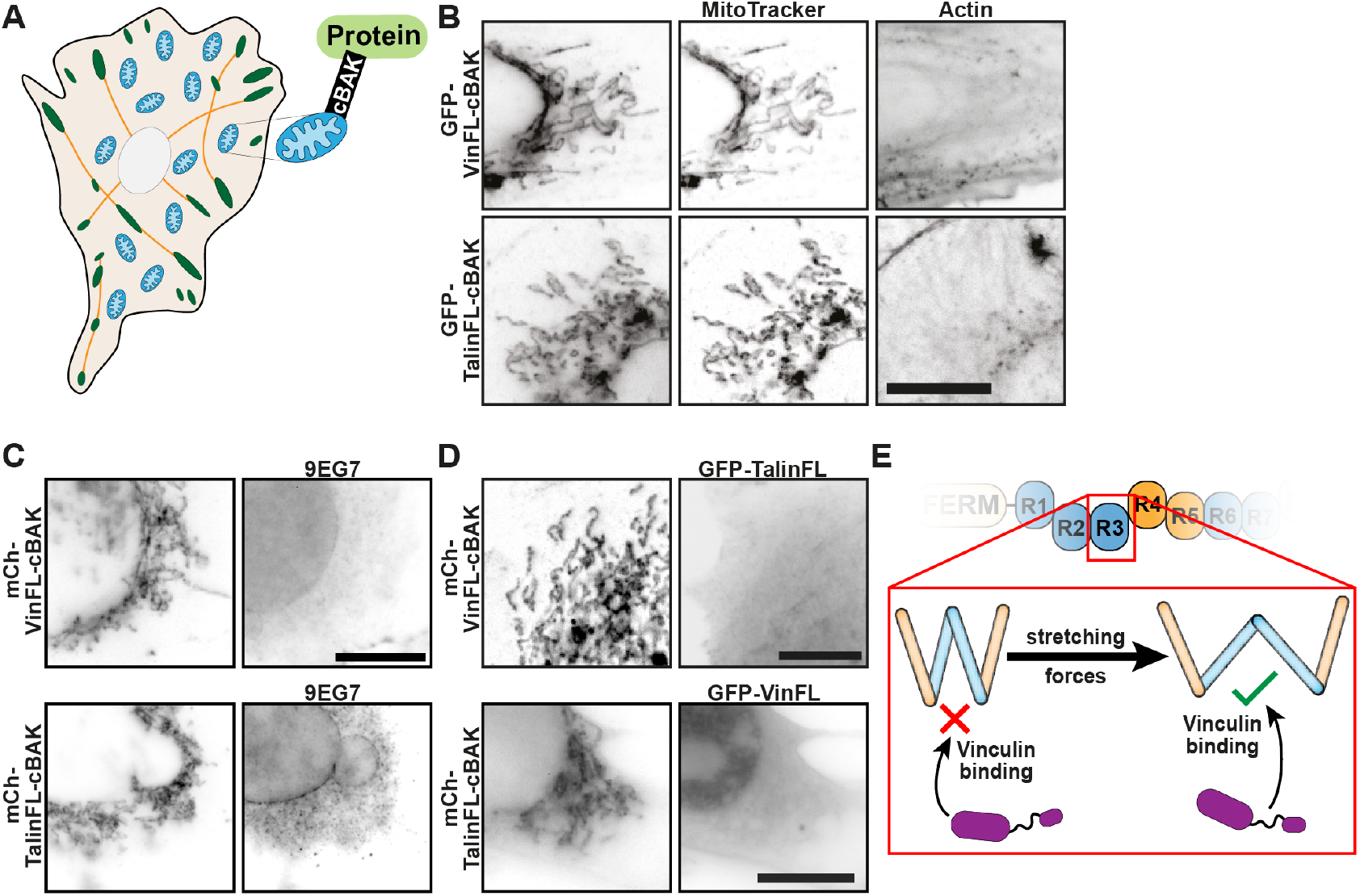
Talin and vinculin do not interact at mitochondria. **A**. The C-terminus of Talin or Vinculin was fused with the short mitochondrial targeting sequence from the outer mitochondrial protein BAK (cBAK). FAs are shown in green, actin stress fibres in orange, mitochondria in blue. **B**. When expressed in NIH3T3 cells, VinFL-cBAK and TalinFL-cBAK both co-localise with the mitochondria-specific dye MitoTracker. Phalloidin staining showed no actin at the mitochondrial surface. **C**. Staining with 9EG7 shows that activated integrins are absent from this system. Scale bars in A, B, C indicate 10 µm. **D**. Co-expression in NIH3T3 cells of GFP-Talin and (mCherry) mCh-vinFL-cBAK, or GFP-VinFL and mCh-TalinFL-cBAK, reveals that full-length vinculin and talin do not interact with each other at mitochondria. **E**. Schematic showing how forces are proposed to be involved in stretching talin rod domains (e.g. R3 4-helix bundle is shown here), exposing previously hidden vinculin binding sites (blue).

### Active vinculin binds talin without forces

These data are in line with *in vitro* single molecule stretching experiments (del Rio et al., 2009; Yao et al., 2014) which show that force acting on helical talin rod domains unmask the cryptic VBS required to support the talin-vinculin interaction (Fig. 1e). Therefore, we hypothesized that in the absence of force talin should not interact with vinculin regardless of the activation state of vinculin. To test this hypothesis, we co-expressed GFP-talinFL with a constitutively active (opened) form of full-length vinculin, vinT12 (Cohen et al., 2005), as well as truncated forms of vinculin (vin258 and vin880) that have exposed talin binding sites, but lack the actin-binding site located in the vinculin tail region (Vt) (Carisey et al., 2013). Each vinculin construct was tagged with cBAK for mitochondrial targeting and mCherry for visualization. Surprisingly, GFP-talinFL bound to all of the vinculin constructs (Fig. 2a, and Supplemental Fig. 1a). Moreover, the interaction occurred in the presence of the actomyosin inhibitors blebbistatin or Y-27632, and also the actin polymerization inhibitor cytochalasin D (Fig. 2b), demonstrating that actomyosin-mediated forces are not essential for talinFL to bind activated vinculin. Similarly, active vinculin at mitochondria also recruited a talinFL construct bearing mutations that compromise the two actin-binding sites (ABS2 and ABS3) in the talin rod (Atherton et al., 2015; Kumar et al., 2016) (Fig. 2c).

**Fig. 2.**
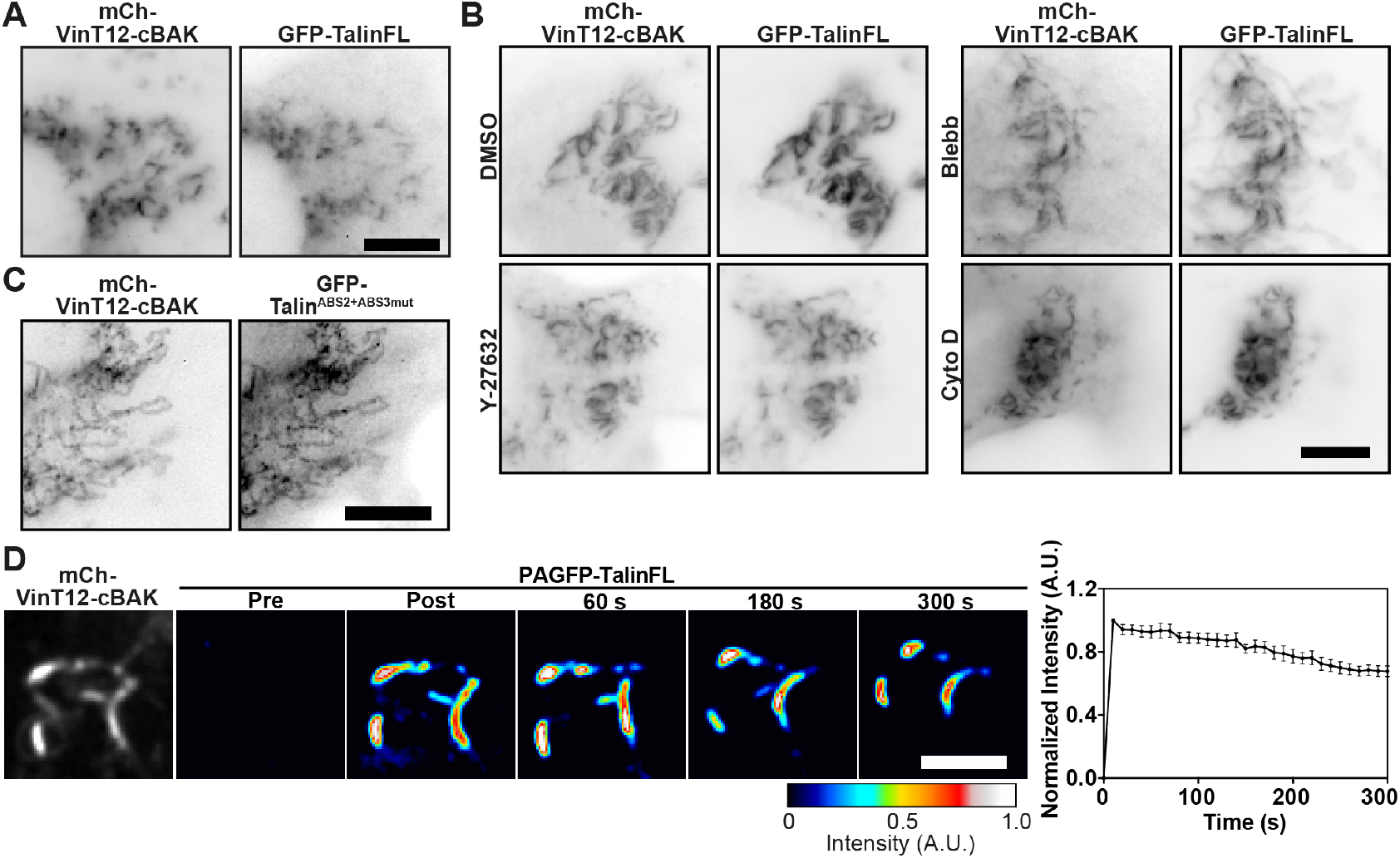
Active vinculin can bind talin independently of forces. **A**. Co-expression of active mCh-VinT12-cBAK with GFP-TalinFL in NIH3T3 cells shows that the two constructs co-localise at mitochondria. **B**. This interaction occurs in the presence of Y-27632 (50 µM) or blebbistatin (50 µM) or Cytochalasin D (Cyto D, 2.5 µg ml-1). **C**. mCh-VinT12-cBAK also recruited a talin construct that has mutations in both central actin binding sites (ABS2 and ABS3, GFP-Talin^ABS2+ABS3mut^) in NIH3T3 cells. Scale bars in A, B, C indicate 10 µm. **D**. Fluorescence Loss After Photoactivation (FLAP) experiments in NIH3T3 cells co-expressing mCh-vinT12-cBAK and Photo-Activatable (PA) GFP-TalinFL show that there is minimal loss of fluorescence over time post activation. Error bars are S.E.M, n = 11 mitochondria from 5 cells. Results are representative of 3 independent experiments. Scale bar indicates 5 µm.

In FAs, increased engagement of talin and vinculin with the actomyosin machinery has been proposed to induce conformations that lead to their activation and thus reduce their mobility (Elosegui-Artola et al., 2016; Humphries et al., 2007). Hence we speculated that the binding of activated/truncated forms of vinculin to talinFL at mitochondria (a site lacking the forces proposed to unmask binding sites in talin and vinculin) might be of low affinity resulting in high turnover rates. However, Fluorescence Loss After Photoactivation (FLAP) experiments revealed that, similar to the turnover of PAGFP-cBAK (Supplemental Fig. 1b, (Schellenberg et al., 2013)) the interaction between talinFL and active vinculin is extraordinarily stable (Fig. 2d) with mobile fractions (Mf) below 20%. Interestingly, the turnover of talinFL bound to vinT12-cBAK at mitochondria was slower than talinFL bound to vinT12 at FAs (Supplemental Fig. 1c). We conclude that vinculin constructs that already have an exposed talin-binding site can activate talinFL and bind to it with high affinity without the involvement of force. This explains the presence of stable force-independent adhesions in the presence of activated vinculin (Atherton et al., 2015; Carisey et al., 2013).

### Active talin disrupts vinculin head-tail auto-inhibition

There are numerous potential VBS throughout the rod ((Gingras et al., 2005), Fig. 3a), and we next aimed to determine which regions of the talin rod can interact with vinculin in the absence of force. To this end, we co-expressed talin constructs lacking domains R4-R10 (GFP-Tal∆R4-R10) or R2-R3 (GFP-Tal∆R2R3) (Fig. 3a) together with constitutively active vinT12-cBAK. Both talin constructs co-localised with vinT12-cBAK at mitochondria (Fig. 3b), demonstrating that VBSs within the N-terminal (R1-R3) and more C-terminal (R4-R13) regions of the talin rod are able to bind activated vinculin independent of force. Surprisingly, these talin deletion constructs also bound to co-expressed wild-type vinFL-BAK (Fig. 3c). A mutation in the talin binding site of vinculin (A50I (Bakolitsa et al., 2004)) blocked the recruitment of vinculin to talin at mitochondria (Supplemental Fig. 2), demonstrating that these talin-vinculin interactions at mitochondria are mediated by the canonical pathway (Bakolitsa et al., 2004). Moreover, FLAP experiments showed that these interactions between vinFL-cBAK and the talin truncation mutants had a similar stability to the interaction between talinFL and vinT12-cBAK (Fig. 3d). From these experiments we conclude that the talin rod contains at least two domains, one in the R2R3 and one in the R4-R10 region that can disrupt the vinculin head-tail interaction, leading to vinculin activation in a force independent manner. The lack of interaction between vinFL and talinFL clearly establishes that the vinculin-activating domains in talin are not accessible in the auto-inhibited talin structure.

**Fig. 3.**
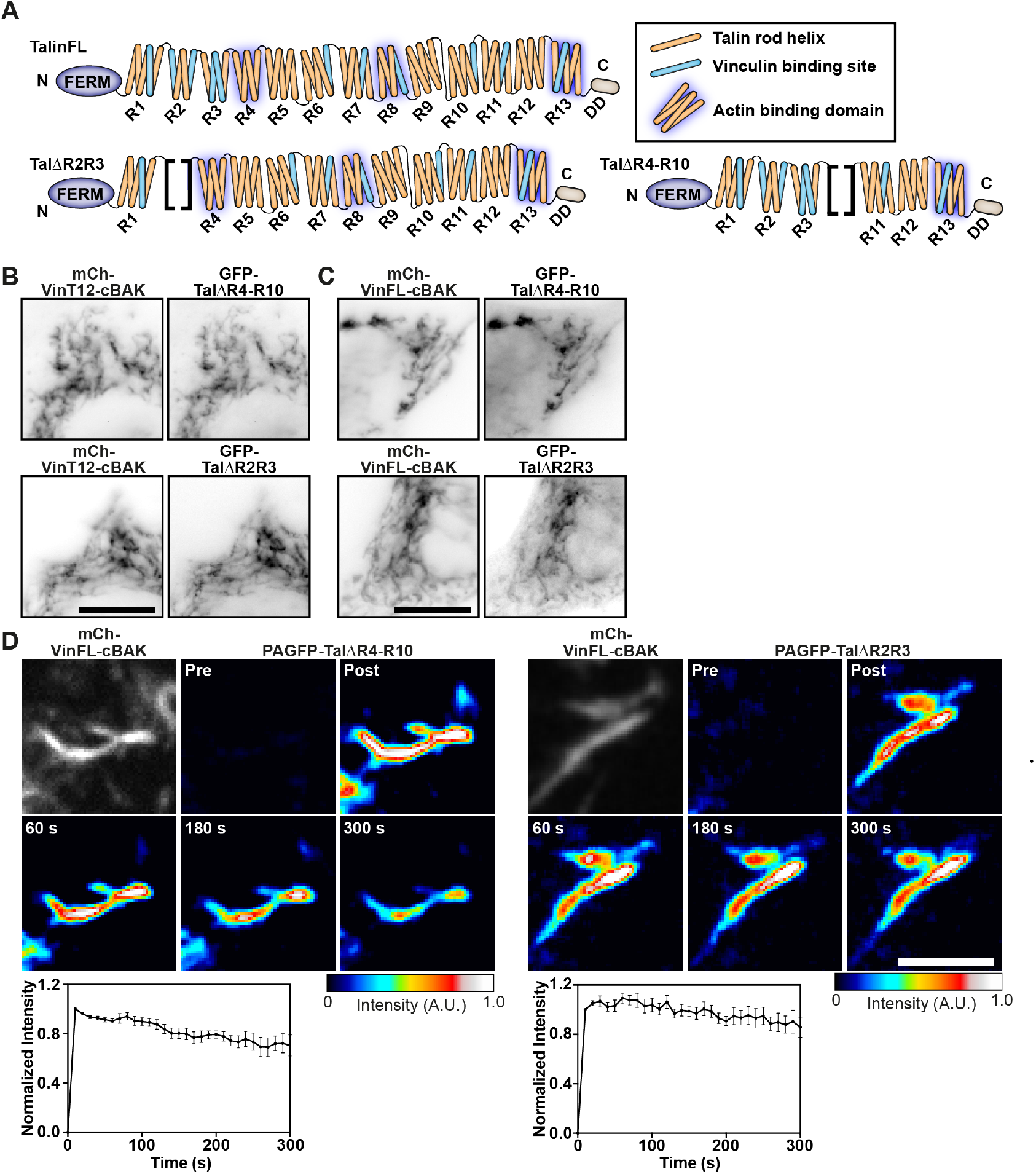
Active talin can bind to vinculin independently of force. **A**. Schematics of the talin constructs used. Blue indicates VBS within the 4-and 5-helix bundles that make up the talin rod (R1-R13). **B, C**. GFP-Tal∆R4-R10 and GFP-tal∆R2R3 are both recruited to either constitutively active mCh-vinT12-cBAK (B) or wild-type mCh-vinFL-cBAK(C) when co-expressed in NIH3T3 cells; scale bar indicates 10 µm. **D**. FLAP experiments in NIH3T3 cells show that there is minimal loss of fluorescence over time, indicating that the interaction between vinFL-cBAK and tal∆R4-R10 (upper panel) and tal∆R2R3 (lower panel) is very stable. Error bars are S.E.M, n = 8 (PAGFP-tal∆R2R3) or 6 (PAGFP-tal∆R4-R10) mitochondria from 5 cells, results are representative of 3 independent experiments. Scale bar indicates 5 µm.

### Relief of talin auto-inhibition is sufficient to induce vinculin binding

The current model of talin (Fig. 4a) suggests the actin and vinculin-binding sites are unavailable in cytoplasmic form of the molecule (Goult et al., 2013a). Previous studies suggested disruption of F3-R9 auto-inhibition as an early step in talin activation (Goksoy et al., 2008), required for integrin binding and activation (Goksoy et al., 2008; Goult et al., 2009). We questioned whether disrupting this F3-R9 interaction would promote the conformational changes required to permit force-independent vinculin binding. To test this hypothesis we introduced an E1770A mutation in the talin R9 domain (talinE1770A) that disrupts the talin F3-R9 interaction, thus relieving talin auto-inhibition (Fig. 4a) (Goult et al., 2009). In contrast to wild-type talin, the talinE1770A mutant readily bound to vinFL-cBAK (Fig. 4b), and FLAP experiments demonstrated that this interaction was very stable (Mf <20%) (Fig. 4c). Similarly, a talin∆FERM rod only construct (Wang et al., 2011) also bound vinFL-cBAK (Supplemental Fig. 3a). These findings demonstrate that disrupting the F3-R9 interaction is sufficient to expose vinculin-activating domains in the talin rod.

**Fig. 4.**
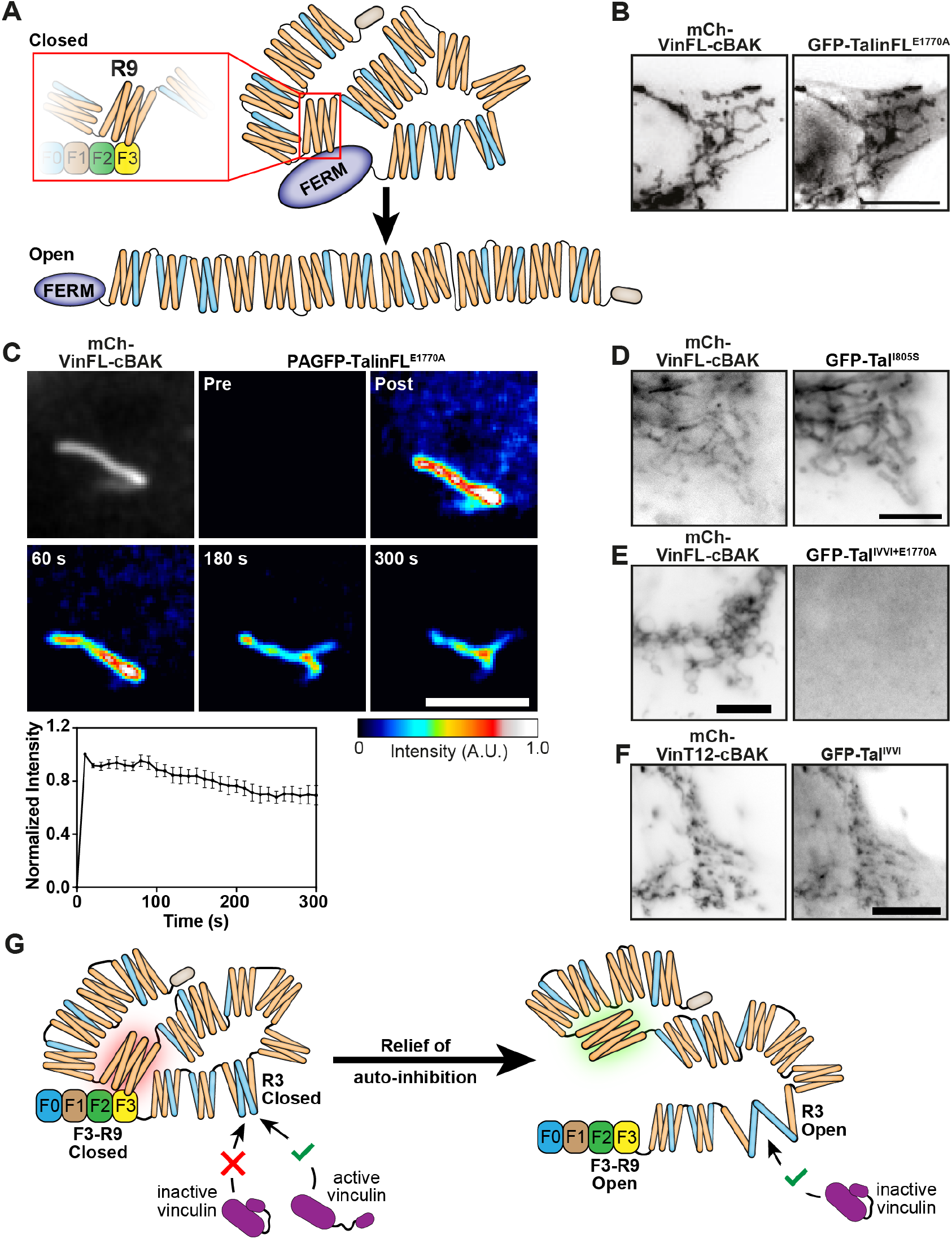
The talin R3 domain is a critical regulator of vinculin binding and talin activation in response to disruption of talin autoinhibition. **A**. Schematic showing talin in a closed compact conformation where the R9 rod domain interacts with F3 of the FERM domain, and an open talin conformation. **B**. GFP-talinFL^E1770A^ (auto-inhibition is compromised) is recruited to mCh-vinFL-cBAK when co-expressed in NIH3T3 cells. Scale bar indicates 10 µm. **C**. FLAP experiments in NIH3T3 cells shows there is minimal loss of fluorescence over time of photoactivated PAGFP-TalinFL^E1770A^ when bound to mCh-vinFL-cBAK at mitochondria. Scale bar indicates 5 µm, error bars show the S.E.M, n = 7 mitochondria from 5 cells. Results are representative of 3 independent experiments. **D**. GFP-TalFL^I805S^ (an R3 destabilising mutant) is recruited even to wild-type mCh-vinFL-cBAK at mitochondria. **E**. A GFP-talin construct in which F3-R9 auto-inhibition is relieved but R3 is stabilised (GFP-Tal^IVVI+E1770A^) is not recruited to wild-type mCh-vinFL-cBAK at mitochondria. **F**. GFP-Tal^IVVI^ (R3 rod domain stabilising mutant) is capable of binding to constitutively active mCh-vinT12-cBAK. Experiments in D, E, F were all performed in NIH3T3 cells, scale bars indicate 10 µm. **G**. Model explaining how relief of talin auto-inhibition regulates the potential for vinculin to bind to the R3 rod domain of talin. In the closed, auto-inhibited talin conformation, only active vinculin is capable of binding to talin R3. When the F3-R9 auto-inhibition is broken, the R3 domain undergoes a conformational change allowing wild-type vinculin to bind.

### The R3 domain of talin is a key determinant of vinculin binding

Vinculin binding to the talin R2/R3 rod domains triggers a conformational change that regulates actin binding to the adjacent R4-R8 domains (ABS2) (Atherton et al., 2015). The experiments above suggest that relief of talin auto-inhibition could induce similar conformational changes in talin permitting vinculin binding. *In vitro* experiments indicate that the talin R3 4-helix bundle, which contains two VBS, is the first helical bundle to unfold in response to force (Yao et al., 2014), and high-pressure NMR experiments show that this domain is inherently unstable, existing in equilibrium between closed and partly open states (Baxter et al., 2017). This is largely due to a cluster of four threonine residues buried within the hydrophobic core of R3. Thus, we hypothesized that introducing additional hydrophilic residues into the R3 core (Tal^I805S^ or Tal^L897S^) that shift the equilibrium towards the open state (Rahikainen et al., 2017) may on their own be sufficient to expose VBS and thereby activate vinculin. Indeed, we found that such mutations triggered binding of mutant talinFL to auto-inhibited vinFL-cBAK (Fig. 4d, Supplemental Fig. 3b).

Conversely, we hypothesized that increasing the stability of R3 would have the opposite effect. Mutating the four threonine residues buried within the R3 core to hydrophobic residues (T809I, T833V, T867V, T901I; GFP-Tal^IVVI^) has previously been shown to stabilize the R3 helical bundle (Yao et al., 2014), requiring more force to stretch this domain (Goult et al., 2013a; Yao et al., 2014). Since talinFL and vinFL do not interact at mitochondria unless one of the two is active, we introduced the IVVI stabilising mutations into the Tal^E1770A^ mutant in which auto-inhibition is relieved.

Remarkably, we detected no interaction between the GFP-TalFL^IVVI-E1770A^ double mutant and vinFL-cBAK (Fig. 4e) in marked contrast to the strong and stable association of the GFP-Talin^E1770A^ mutant with vinFL-cBAK at mitochondria. This result is important since it suggests that, following relief of talin auto-inhibition, rearrangements in the R3 domain are absolutely critical for binding to vinFL, and the exposure of the talin-binding site in the vinculin head. However, to our surprise, a GFP-Tal^IVVI^ construct was recruited to activated vinT12-cBAK (Fig. 4f) at mitochondria, as well as to the vin258-cBAK construct (vinD1 domain only) (not shown). These results show that active vinculin can even bind to and activate auto-inhibited talin containing a stabilised R3 domain.

In summary, these results lead to the following model (Fig. 4g). Initially, activated talin (after relief of auto-inhibition) binds to vinFL via the VBS within the talin R3 helix. Binding to this VBS is sufficient to activate vinculin by disrupting the vinculin head-tail interaction, thus facilitating further vinculin binding to talin. Once activated, vinculin (with an exposed talin-binding site) can then bind to talinFL at multiple VBS within the talin rod independent of force.

### Talin, vinculin and force-independent FA assembly

Given the surprising finding that talin and vinculin can interact at mitochondria in the absence of force, we sought to clarify the role of actomyosin-mediated tension during FA formation and maturation. Tension release inhibits maturation of adhesion complexes into streak-like FAs, although small adhesions complexes containing talin and vinculin still remain (Stutchbury et al., 2017). To mimic experiments at mitochondria, we co-expressed activated mCh-vinT12 together with auto-inhibited GFP-talinFL in TalinKO cells, and compared FA formation to cells co-expressing auto-inhibited mCh-vinFL and GFP-talinFL. We created a force-free environment by pretreating cells with the tension-releasing drug blebbistatin (50 µM) for 45 minutes, before allowing them to spread on fibronectin for 15 minutes. Cells co-expressing auto-inhibited mCh-vinFL and GFP-talinFL only formed small peripheral adhesions (Fig. 5A). In contrast, cells co-expressing mCh-vinT12 and GFP-talinFL formed larger adhesions throughout the cell that were positive for both talin and constitutively active vinculin (mCh-vinT12) (Fig. 5B). Interestingly, although these adhesions were streak-like as seen in mature FAs of non-treated cells, they were random in orientation and often bent (Fig. 5C, Supplemental Fig. 4A). We wondered whether this buckling was due to the lack of tension resulting from inhibition of the actomyosin-machinery. To test this, we treated vinKO MEFs expressing GFP-vinT12 and RFP-LifeAct with Y-27632 (50 µM). Remarkably, the previously well-organised and streak-like FAs with vinT12 attached to actin stress fibers started to bend and buckle without being disassembled (Fig. 5D, Supplemental Movie 1). Importantly, at the cell periphery, new adhesions formed that seemingly grow even under tension-release conditions (Fig. 5E, Supplemental Movie 1). These experiments suggest that in FAs, activated vinculin can activate talin and link the complex to actin filaments. Whilst there seems to be some bundling activity through the binding of talin and vinculin, full maturation into a stress fiber-associated tensile FA requires actomyosin-mediated tension.

**Fig. 5.**
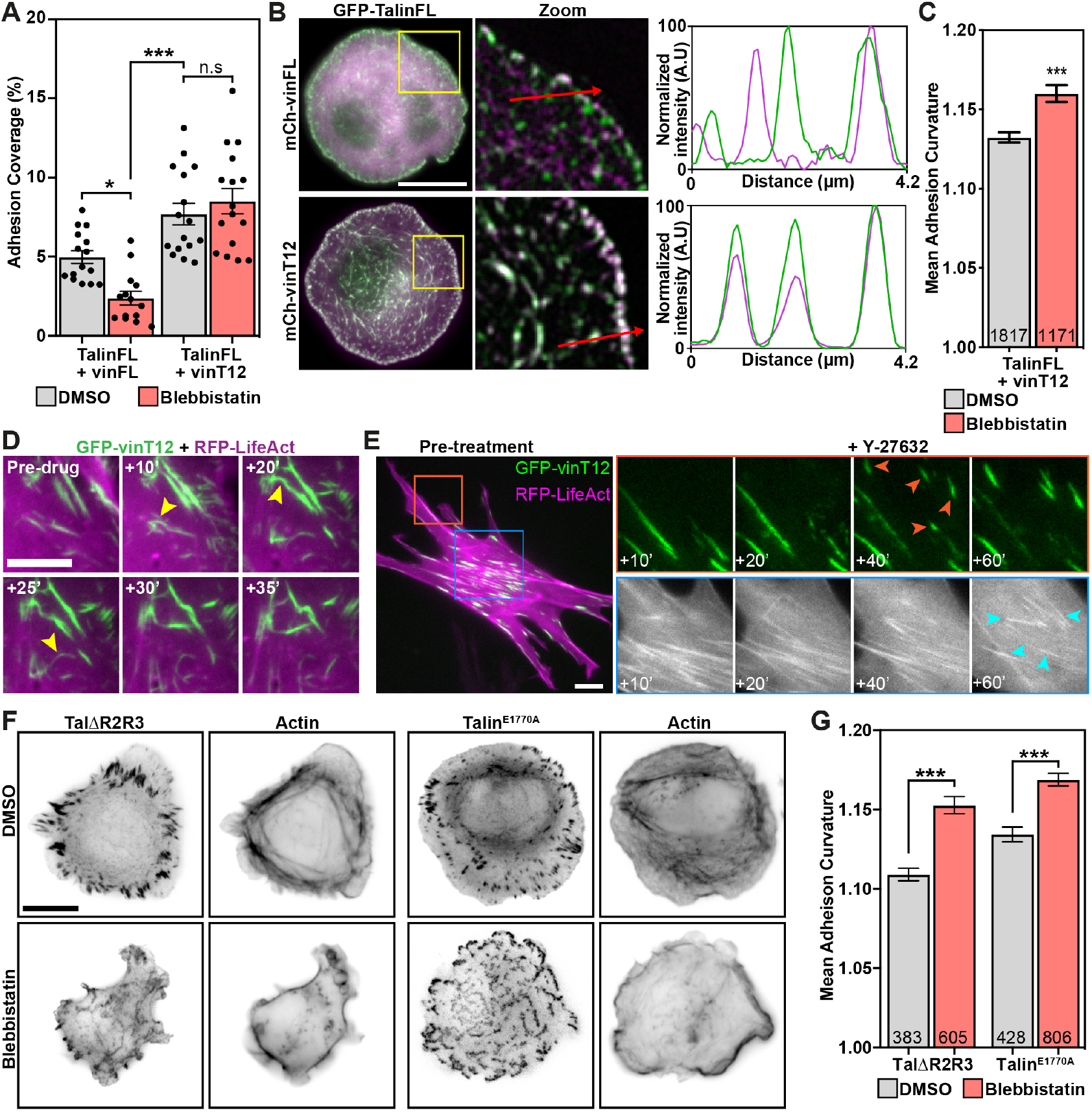
Active vinculin or active talin support adhesion formation under tension-release conditions. **A**. TalinKO cells co-expressing GFP-TalinFL with either auto-inhibited mCh-vinFL (left) or activated mCh-vinT12 (right) were pre-treated in suspension with blebbistatin (50 µM) or an equivalent volume of DMSO, for 45 minutes. Cells were fixed after spreading on fibronectin-coated glass for 15 minutes. The percentage of the cell consisting of adhesions was quantified from the GFP signal. Graphs show the mean and S.E.M; results are representative of 3 independent experiments. **B**. Representative images of talinKO cells as described above. Magnified region (yellow box) has been background subtracted and smoothened. Line profiles drawn in the direction indicated by the red arrow show that mCh-vinFL and GFP-vinFL only co-localise at peripheral adhesion structures, whereas mCh-vinT12 and GFP-vinFL co-localise at all adhesions throughout the cell. **C**. Quantification of the curvature of talin-positive structures in these cells. Graphs show the mean and S.E.M; values in the bars indicate the numbers of structures analysed from 16 (DMSO) and 15 (Blebbistatin) cells. Results are representative of 3 independent experiments. **D**. Still frame images from a movie of a vinculinKO MEF expressing GFP-vinT12 (green) and RFP-LifeAct (magenta) before and during treatment with Y-27632 (50 µM). Note the buckling and bending of the adhesions (yellow arrows). Scale bar indicates 5 µm. **E**. VinculinKO MEF expressing GFP-vinculinT12 together with RFP-LifeAct treated with Y-27632 (50 µM) and imaged every minute for 60 minutes. Active vinculin supports the formation and growth of new adhesions (upper panels, orange arrows), with actin stress fibres remaining bundled at the centre of the cell (lower panels, cyan arrows). **F**. TalinKO cells expressing GFP-Tal∆R2R3 or GFP-Talin^E1770A^, after DMSO or blebbistatin treatment (as described above in A), with phalloidin staining shows GFP-positive structures form during cell spreading without intracellular tension. **G**. Quantification of the curvature of talin-positive structures in these cells. Graph shows the mean and S.E.M; values in the bars indicate the numbers of structures analysed from 15 cells for each condition. Results are representative of 3 independent experiments. Scale bars in B, E, F indicate 10 µm. In A, C, G, * indicates p<0.05; *** indicates p<0.001.

We next investigated whether active talin constructs could support adhesion maturation in the absence of forces (after treatment in suspension with blebbistatin as described above). As for vinT12, expressing either activated tal∆R2R3 or talin^E1770A^ in TalinKO cells promoted the formation of talin-positive elongated and disorganized FA structures that were larger than those found in control cells expressing talinFL (Fig. 5F, G, Supplemental Fig. 4B, C).

### Paxillin can bind to both talin and vinculin independently of force

Talin and vinculin orchestrate adhesion signaling by binding to many other proteins within the FA including paxillin, which binds to both talin and vinculin (Turner et al., 1990; Wood et al., 1994; Zacharchenko et al., 2016). Additionally, paxillin is thought to contribute to the recruitment of vinculin to focal complexes (Case et al., 2015; Pasapera et al., 2010). To examine whether forces are required for the association of paxillin with talin and vinculin, we probed their interaction using the mitochondrial targeting assay. Interestingly, both GFP-Paxillin and endogenous paxillin were recruited to both vinFL-cBAK and talinFL-BAK (Fig. 6a, Supplemental Fig. 5A). Since the wild-type forms of talin and vinculin don’t interact with each other at mitochondria we conclude that they can both associate with paxillin in their inactive form. Similarly, a mCh-Paxillin-cBAK construct was able to recruit both GFP-Talin and GFP-Vinculin to mitochondria (Supplemental Fig. 5B).

**Fig. 6.**
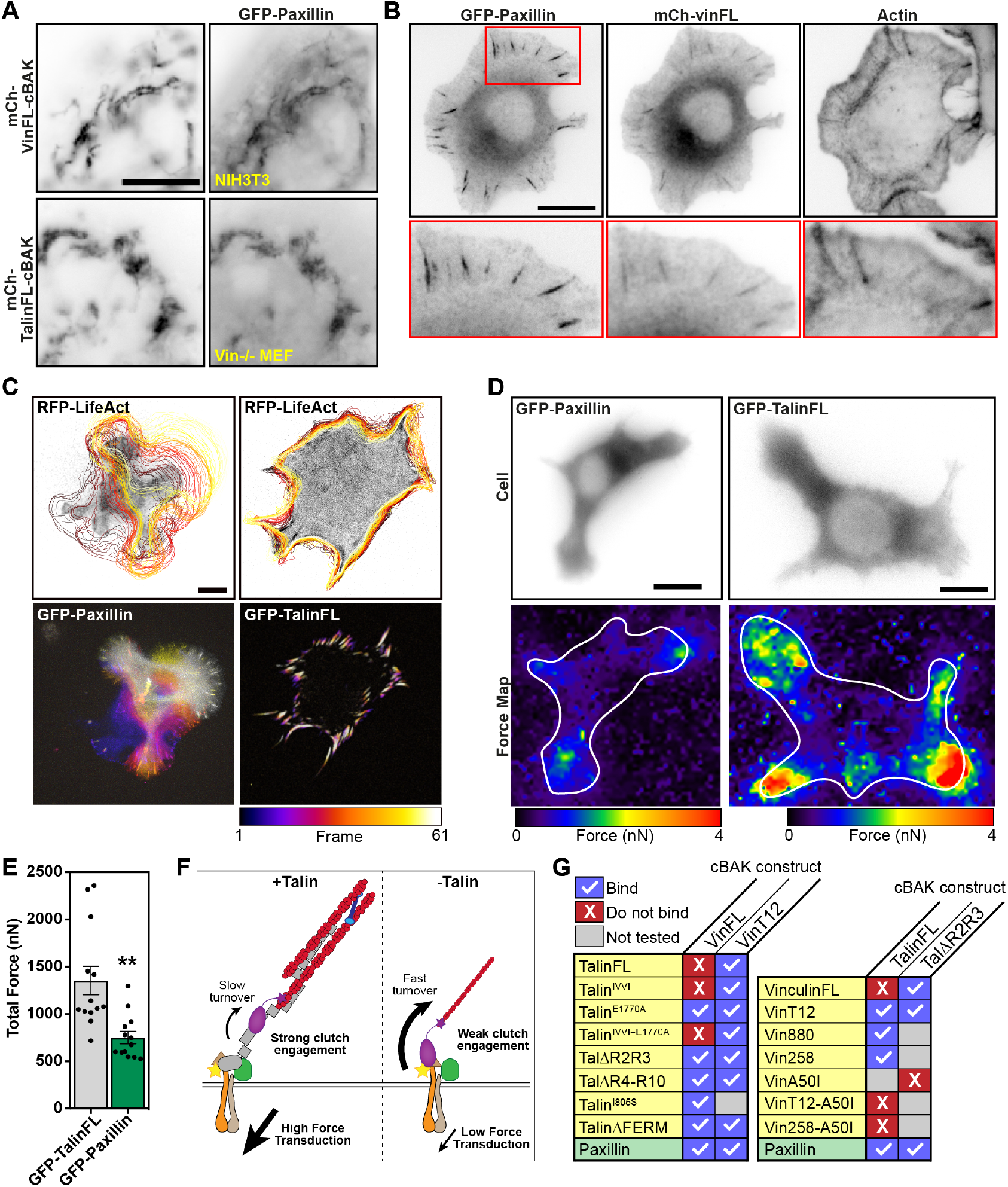
Paxillin can bind both talin and vinculin when either is in an inactive conformation, independently of forces. **A**. Co-expression of GFP-Paxillin with either mCh-vinFL-cBAK or mCh-TalinFL-cBAK in NIH3T3 cells reveals that paxillin can bind to inactive vinculin or talin independently of forces. **B**. TalinKO cells expressing GFP-Paxillin and mCh-vinFL were fixed after spreading on fibronectin for one hour in the presence of manganese (5 mM) and stained with 647-Phalloidin. **C**. Live-cell imaging of TalinKO cells co-expressing either GFP-Paxillin or GFP-TalinFL with RFP-LifeAct (top panel). The cell edge was traced over time using the RFP-LifeAct signal. Black outline indicates the cell position at the first frame; yellow indicates the position in the last frame. Temporal colour maps of adhesion movement obtained from the GFP signal of the same cell (lower panel) show that cells without talin after Mn^2+^ treatment are highly dynamic compared to talin-expressing Mn^2+^-treated cells. Images were acquired every 2 minutes for 2 hours. **D**. Representative force maps of talinKO cells expressing either GFP-Paxillin or GFP-TalinFL spread on a polyacrylamide hydrogel containing fluorescent beads for traction force microscopy. White line indicates the outline of the cell. Blue colour indicates regions of low force exertion; red indicates regions of high force exertion. Scale bars in A-D indicate 10 µm. **E**. Quantification of the total force exerted per cell from traction force microscopy experiments. Graphs show the mean and S.E.M; n = 13 cells, results are representative of 3 independent experiments; ** indicates p<0.01 (t-test). **F**. Schematic showing the role of talin in force transduction. In wild-type cells (+talin), vinculin (purple) reinforces the link between talin (grey) and actin, engaging the molecular clutch for efficient force transduction, stabilising adhesion turnover. Without talin (-talin), vinculin is weakly anchored to additional adhesion proteins, and is only able to transduce low forces. These adhesions are unstable and rapidly turned-over. **G**. Summary tables showing which talin or paxillin constructs bind to which vinculin-cBAK constructs (left) or which vinculin or paxillin constructs bind to which talin-cBAK constructs (right).

To examine the potential role of paxillin in recruiting vinculin to adhesions in the absence of talin, we conducted experiments in talinKO cells. Under normal conditions these cells neither adhere nor spread, but activation of integrins by Mn2+ induces spreading on fibronectin (Theodosiou et al., 2016). Surprisingly, in Mn2+-stimulated talinKO cells, we found that both paxillin and vinculin localized to FA-like structures located along filopodia-like actin bundles embedded in protruding lamellipodia (Fig. 6b, Supplemental Fig. 5c). Live-cell imaging of talinKO cells expressing GFP-Paxillin and RFP-LifeAct stimulated with Mn^2+^ revealed that the actin cytoskeleton of these cells was extremely dynamic, and that the adhesions had a fast turnover. This was in stark contrast to talinKO cells expressing GFP-TalinFL and RFP-LifeAct stimulated with Mn^2+^, which displayed very stable adhesions (Fig. 6c, Supplemental Movie 2). Furthermore, traction force microscopy in presence of Mn^2+^ revealed that FAs in talinKO cells generated only about 50% of traction forces compared to control cells expressing GFP-TalinFL (Fig. 6d, e).

Taken together, these experiments demonstrate that talin is not essential for the recruitment of either paxillin or vinculin to FA-like structures if cell spreading is induced by the artificial activation of integrins with Mn^2+^. Moreover, together with the data derived from the mitochondrial targeting system, the data above raise the possibility that paxillin and vinculin can support the mutual recruitment of each other to adhesions even when vinculin is in an inactive state.

## Discussion

The development of FAs is a highly dynamic process where growth is associated with increasing molecular complexity and actomyosin contractility. The interaction between talin and vinculin is thought to play a key role in the assembly of nascent adhesions, but the short lifetime of these structures makes investigating the early events in adhesion formation difficult. Moreover, the complex nature of protein-protein interactions in mature FAs makes studying such interactions problematic. To overcome these barriers, we used a mitochondrial-targeting assay to examine molecular interactions at mitochondria, which move in a force-free environment in the cell’s cytoplasm independent of actomyosin. We modified the assay used previously (Bubeck et al., 1997; Cohen et al., 2006; Maartens et al., 2016) by employing the C-terminus of BAK (cBAK) instead of ActA as the targeting motif. In contrast to ActA, the cBAK motif stably integrates into the outer mitochondrial membrane (Supplemental Fig. 1b, (Schellenberg et al., 2013)), allowing the assessment of protein-protein binding strength using FLAP, and nullifying movement of proteins between FAs and mitochondria. Our results provide important new insights into the molecular mechanisms underpinning the talin/vinculin interaction, and the assembly of FAs.

*In vitro* stretching experiments show that force exerted on the helical bundles of the talin rod exposes the multiple VBSs contained therein (del Rio et al., 2009; Yao et al., 2014; Yao et al., 2016), suggesting force across talin is required for the vinculin/talin interaction. However, the results from our force-free mitochondrial targeting system do not fit this model. While as expected, auto-inhibited vinFL-cBAK does not bind auto-inhibited talinFL-cBAK, (Fig. 1d), we found that (i) activated vinculin constructs stably interacted with talinFL-cBAK (Fig. 2a, Supplemental Fig. 1a) and (ii) a talin construct lacking the FERM domain bound to vinFL-cBAK (Supplemental Fig. 3a). Moreover, experiments with talin deletion constructs (tal∆R2R3, tal∆R4-R10) that also bind to vinFL-BAK, suggest that there are multiple VBS within the talin rod that are accessible to vinculin without the need of forces. Indeed, none of the above interactions were affected by blebbistatin, Y-27632 or cytochalasin D demonstrating that force exerted by actomyosin contraction is not essential for activated vinculin to bind talinFL or vice versa. Similar stable interactions between activated vinculin constructs and full-length talin have been observed in the cytoplasm of Drosophila embryos, (Maartens et al., 2016), a site where forces would not be expected to contribute to complex assembly.

How can one explain the discrepancy between previous “force-induced” models and our data? Careful inspection of *in vitro* single molecule experiments show that there is some residual binding of vinD1 to unstretched talin rod domains (del Rio et al., 2009; Goult et al., 2013b), and perhaps the cellular environment favours more labile conformations of the talin helices that support vinculin binding. A key factor may be temperature; *in vitro* stretching experiments were conducted at room temperature (Yao et al., 2014) whereas the cellular milieu is 37°C. Such speculation is in line with data on the interaction between vinD1 and the talin R1 (Patel et al., 2006), R2 and R3 (Goult et al., 2013b) rod domains. Whilst R3 bound vinD1 at 25°C, the other two domains required the temperature to be increased to 37°C before binding was observed. The different levels of thermal energy required to induce complex formation are also in line with the different levels of mechanical energy required to unfold the different talin rod domains. Thus, R3 is the least stable, and unfolds at 5 pN whereas R1 and R2 unfold at 15-24 pN (Yao et al., 2016). Even R3 containing the IVVI stabilizing mutation unfolds at 8 pN (Yao et al., 2014), a force well below that required to unfold R1/R2. Therefore, it is not surprising that vinT12-cBAK (and vin258-cBAK, data not shown) can bind talinFL containing the IVVI mutation (Fig. 4f).

The finding that a talin rod-only construct (talin∆FERM) unlike talinFL binds to vinFL-cBAK (Supplemental Fig. 3a) indicates that the FERM domain plays a critical role in regulating the accessibility of the VBS in the talin rod. Indeed, the F3 region of the talin FERM domain is known to interact with the R9 domain of the rod to provide auto-inhibition (Goult et al., 2013a) (Fig. 4a). Our data showing that relief of auto-inhibition alone (using an E1770A point mutation in R9 (Goksoy et al., 2008; Goult et al., 2009; Haage et al., 2018)) is sufficient to induce binding of talinFL to vinFL-cBAK (Fig. 4b) clearly demonstrates that disrupting the F3-R9 interaction unmasks VBS in the talin rod.

Significantly, talinFL containing the E1770A mutation plus the R3 domain-stabilizing IVVI mutations (tal^IVVI-E1770A^) was unable to bind to vinFL-cBAK, suggesting a model whereby release of talin auto-inhibition leads to a conformational change in R3 that facilitates vinculin binding (Fig. 4g). The finding that point mutations in R3 that increase the lability of this domain (tal^I805S^ and tal^L897S^ (Rahikainen et al., 2017)), resulted in binding of talinFL to vinFL-cBAK without the need for mutations that relieve auto-inhibition support the idea that talin auto-inhibition and the stability of R3 go hand in hand.

Precisely how talin auto-inhibition is regulated *in vivo* remains to be determined. One proposed mechanism is the “pull-push” hypothesis (Song et al., 2012) in which plasma membrane localized PIP_2_ and Rap1, plus Rap1-binding partners RIAM and lamellipodin (all of which interact with talin), are envisaged to co-operate to expose the talin F3 domain promoting binding to the cytoplasmic tail of β-integrin (Goult et al., 2013b; Lagarrigue et al., 2015; Lee et al., 2009; Song et al., 2012; Yang et al., 2014). Interestingly, a talin construct containing mutations in the FERM domain that block the interaction between talin and PIP_2_ was unable to support integrin activation (Goult et al., 2010) or rescue FA formation in talin-null cells (Chinthalapudi et al., 2018), suggesting that lipid-binding is critical for the recruitment of talin to the plasma membrane prior to activation. PIP_2_ has also been reported to disrupt the vinculin head-tail interaction (Chinthalapudi et al., 2014; Gilmore and Burridge, 1996; Johnson et al., 1998), which our results show enables vinculin to bind to and activate talinFL (Fig. 2).

Since a number of signalling pathways have the potential to trigger talin/vinculin activation, is there any role for forces in this process? The idea that forces may contribute to the relief of talin auto-inhibition is consistent with our observations that FA assembly is perturbed in 50% of talinKO cells re-expressing a talin construct bearing mutations in the C-terminal ABS3 domain that inhibits engagement with the actomyosin machinery (Atherton et al., 2015). Besides contributing to relief of talin auto-inhibition, stretching could also promote vinculin recruitment to some of the more stable VBS-containing talin domains. For vinculin, force-mediated activation seems less likely in light of our results: exertion of force across vinculin would require actin binding at the C-terminus and talin binding at the N-terminus, sites that are buried in the crystal structure of auto-inhibited vinculin (Cohen et al., 2006; Izard et al., 2004). Our data show that activation of talin and binding to vinculin through the canonical binding site in vinD1 is sufficient to activate vinculin (Fig. 3c, Supplemental Fig. 2d) implying vinculin auto-inhibition can be relieved independent of force. This is supported by structural studies showing talin is capable of displacing the vinculin tail from the head (Izard et al., 2004).

Integrin clustering can occur independently of intracellular contractility and is enhanced by the presence of talin (Changede et al., 2015) and kindlin2 (Sun et al., 2019; Theodosiou et al., 2016). Our results show that myosin-mediated forces are neither required for talin or vinculin activation, nor for initial adhesion formation (Fig. 5a, b; (Stutchbury et al., 2017)). Rather, myosin-derived forces are required to organize focal complexes into a polarized FA: myosin inhibition during cell spreading did not prevent adhesion formation in cells expressing active vinculin (vinT12, Fig. 5b) or active talin (talin^E1770A^, Fig. 5f), although these proteins clustered into structures with a random orientation (Fig. 5c, g). Therefore, one of the critical functions of myosin is to generate retrograde flow and the bundling of actin filaments linked to adhesion complexes. This drives the coalescence of small adhesions into larger more efficient force-transducing focal contact points as has been proposed previously (Ballestrem et al., 2001; Oakes et al., 2012).

One might wonder why the talin^E1770A^ or vinT12 FA complexes remain so stable without the activity of myosin-II (Fig. 5)? One possibility is that integrins are constitutively activated by talinE1770A and therefore turnover is blocked. However this is not in line with the observations in Mn^2+^-treated talinKO cells where integrins are chemically activated, but adhesions are rather short-lived (Fig. 6c, Supplemental Movie 2). Another possibility is that constitutive binding of activated talin to integrins might block access of other proteins such as SNX17 that binds to the same NPxY motif of the cytoplasmic domain of the beta1 integrin chain (Bottcher et al., 2012). SNX17 is involved in endocytic trafficking of transmembrane proteins, and constitutive talin binding to integrins could therefore prevent integrin endocytosis and recycling, and stabilise adhesion complexes.

The adapter protein paxillin is a critical regulator of cell migration (Deakin and Turner, 2008) and is present at the early stages of adhesion complex formation (Choi et al., 2008; Zaidel-Bar et al., 2007). Paxillin can bind to numerous proteins including kindlin2 (Bottcher et al., 2017), talin (Zacharchenko et al., 2016), vinculin (Deakin et al., 2012; Turner et al., 1990) and FAK (Brown et al., 1996; Turner and Miller, 1994) that are all implicated in integrin activation and cell migration (Deakin and Turner, 2008). Interestingly, our mitochondrial-targeting assay revealed that paxillin binds to both talin and vinculin independent of their activation state and force (Fig. 6a, Supplemental Fig. 5b). Binding is also independent of phosphotyrosine phosphorylation because staining for phosphotyrosine was negative at mitochondria (Supplemental Fig. 5d).

Force-independent binding of paxillin to talin and vinculin raises the possibility that paxillin might participate in the recruitment of both vinculin (as suggested previously (Case et al., 2015; Pasapera et al., 2010)) and talin, possibly in their auto-inhibited forms, to early sites of cell-ECM adhesion in migrating cells. This would be in line with observations that moderate knock-down of both paxillin and Hic5 (a paxillin family member) together reduced the recruitment of talin and vinculin to adhesions, and complete paxillin/Hic5 knockdown prevented the formation of discrete substrate adhesions (Pasapera et al., 2010). Given that cells without talin neither adhere nor spread (Atherton et al., 2015; Theodosiou et al., 2016; Zhang et al., 2008), one might wonder which comes first in the hierarchy of adhesion complex assembly? One possibility is that talin is critical for integrin engagement at the initial attachment phase when a cell encounters ECM. However, as soon as integrins are engaged and actin polymerization initiates cell spreading, adhesion formation can occur independently of talin (Fig. 6).

In wild-type cells, the actin polymerization required for cell spreading downstream of talin-integrin binding can be driven by the recruitment of ENA-VASP proteins to a complex containing activated integrins, talin, and Mig-10/RIAM/lamellipodin (MRL) family proteins (Lagarrigue et al., 2015), with polymerizing actin at the leading edge acting to position conformationally-active integrins (Galbraith et al., 2007). Adding Mn2+ to the culture medium can initiate cell spreading in TalinKO cells (Fig. 6, (Theodosiou et al., 2016; Zhang et al., 2008)), suggesting there are alternative, talin-independent pathways for actin polymerization following integrin activation. In Mn^2+^-stimulated TalinKO cells, both paxillin and vinculin co-localised in these rather short-lived adhesion structures located in the protruding cell periphery at the base of very dynamic filopodia (Fig. 6b, Supplemental Movie 2), which in contrast to talin containing control cells, were not connected to stress fibers and not able to produce larger traction forces (Fig. 6c, d, e). Therefore, talin appears to be an essential component of the molecular clutch acting as a catch bond; the presence of vinculin anchored to additional FA components alone is not sufficient for effective force transduction (Fig. 6f).

Based on our results and previous literature we propose the following model of adhesion formation during polarized cell migration (Fig. 7): at the leading edge, Rap1 and lamellipodin/RIAM increase actin polymerization (Lagarrigue et al., 2015). These proteins facilitate polymerization through actin binding proteins (e.g. Arp2/3, VASP; (Lafuente et al., 2004)), which in turn contribute to the de-novo positioning of proteins such as FAK (Serrels et al., 2007; Swaminathan et al., 2016), kindlin and paxillin (Bottcher et al., 2017) to form pre-adhesion complexes. Together, they can then recruit vinculin (Case et al., 2015; DeMali et al., 2002; Pasapera et al., 2010) and talin (Lawson et al., 2012; Theodosiou et al., 2016); we suggest in their inactive conformations. Once talin auto-inhibition is relieved (potentially through PIP_2_ (Song et al., 2012), and possibly FAK (Lawson et al., 2012)), talin F3 binds β-integrins and ABS3 binds actin. Relief of the F3/R9 interaction also allows the R3 domain to undergo a conformational change exposing its two VBS. Vinculin binding to R3 then unlocks the central ABS2 of talin, and promotes vinculin head-tail dissociation, allowing the vinculin tail to reinforce the link to actin (Atherton et al., 2015). The thereby engaged “molecular clutch” (Thievessen et al., 2013) stabilizes adhesions, with contractile forces increasing clustering of adhesion complexes to mature into larger focal contacts that efficiently mediate traction forces and mechanosensing.

**Fig. 7.**
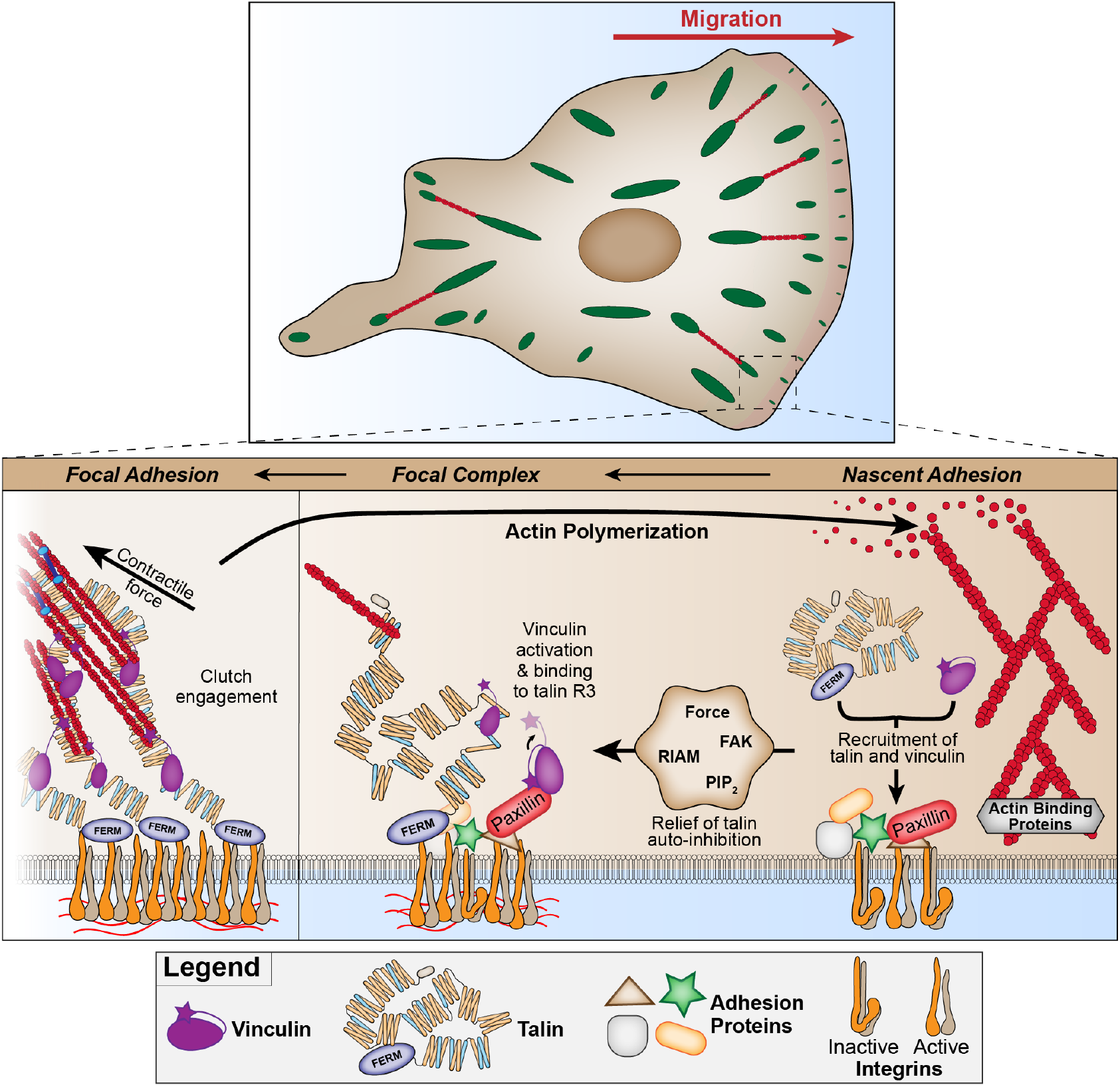
Model of early events in adhesion formation during polarised migration. Actin binding proteins (e.g. Arp2/3, VASP) at the leading edge of a migrating cell trigger the de novo recruitment of proteins (e.g. FAK, kindlin, RIAM and paxillin) to integrins. These proteins lead to the recruitment of talin and vinculin through direct or indirect interactions. Relief of talin auto-inhibition occurs through one of several mechanisms (PIP_2_, FAK, RIAM, or force from actin binding to the C-terminal ABS3), producing a change in conformation within the R3 domain that exposes vinculin binding sites. Vinculin is ‘handed-over’ from paxillin to talin. Vinculin binding to talin R3 promotes further conformational changes, exposing the central ABS2 and engaging the molecular clutch to control force transmission and adhesion maturation.

## ACKNOWLEDGEMENTS

C.B. acknowledges the Biotechnology and Biological Sciences Research Council (BBSRC) and Wellcome Trust for funding of this project. The CB laboratory is part of the Wellcome Trust Centre for Cell-Matrix Research, University of Manchester, which is supported by core funding from the Wellcome Trust (grant number 203128/Z/16/Z). P.A. is funded by BBSRC (BB/P000681/1). QuimP (Baniukiewicz et al., 2018) was developed at the University of Warwick with support from BBSRC (BBR grant BB/M01150X/1). The Bioimaging Facility microscopes were purchased with grants from BBSRC, Wellcome Trust, and the University of Manchester Strategic Fund. We thank the staff of the Bioimaging facility (in particular Dr Peter March) at the University of Manchester for their help with imaging and analysis, Dr M. Ptushkina for generation of constructs, and Dr D. Jethwa for generation of pilot data.

## AUTHOR CONTRIBUTIONS

P.A. and C.B designed experiments, with input from I.B., and D.C. P.A. performed the experiments and analysed the data. F.L. provided assistance in performing experiments. A.C. provided movies used in Figure 5. A.G. provided assistance with generation of cBAK constructs. C.B. conceived ideas with input from P.A. and I.B. P.A. made the figures; P.A., D.C., and C.B. wrote the manuscript with input from A.C. and I.B. C.B. supervised the project and acquired funding.

## Materials and Methods

### Cell Culture

NIH3T3s and vinculin-/-MEFs were cultured in Dulbecco’s modified Eagles medium (DMEM) supplemented with 10% FCS (Lonza), 1% L-glutamine (Sigma), and 1% Non-essential amino acids (Sigma). Talin1/2 double null cells (Atherton et al., 2015) were cultured in DMEM:F12 (Lonza) supplemented with 10% FCS (Lonza), 1% L-glutamine (Sigma), 15 µM HEPES (Sigma) and 1% Non-essential amino acids (Sigma). Transient transfections were performed using Lipofectamine and Lipofectamine Plus reagents (Invitrogen), as per the manufacturer’s instructions. For live-cell imaging and fixed cell imaging, cells were cultured on glass-bottom dishes (IBL, Germany) coated with bovine fibronectin (Sigma) at a final concentration of 10 µg ml-1.

### Generation of cBAK-tagged constructs

To generate vinculin-cBAK constructs, assembly PCR was first used to generate the 108 bp mitochondrial targeting sequence from the C-terminal tail of BAK. The following 4 primers were used: 1) Forward external: TATGAATTCTTGCGTAGAGACC-CCATCCTG, 2) Forward internal: CCCATCCTGACCGTAATGGTGATTTTTGGT, 3) Reverse internal: ATCTGTGTACCACGAATTGGCCCAACAGAA, 4) Reverse external: TATGGTACCTCATGATCTGAAGAATCTGTG. The 5’ and 3’ end primers (external) contained EcoR1 and Kpn1 restriction digestion sites respectively. Site-directed mutagenesis was used to remove the stop codon from vinculin constructs and to add an EcoR1 restriction site. Digestion with EcoR1 and Kpn1 FastDigest enzymes (Fermentas) was used to clone the cBAK fragment into the vinculin (gallus gallus) constructs in Clontech C1 vectors. To generate talin-cBAK constructs, a 1133 bp sequence was synthesised (Genewiz Inc.), consisting of 1031 bp from talin1 (mus musculus) joined to the cBAK fragment (TTGCGTAGAGACCCCATCCT-GACCGTAATGGTGATTTTT GGTGTGGTTCTGTTGGGCCAATTCGTGGTACACA-GATTCTTCAGATCATGA) with the talin stop codon removed, flanked by a 5’ SalI restriction site and a 3’ SacII restriction site. This fragment was cloned into the GFP- and mCherry-talin constructs in Clontech C1 vectors by restriction digest, using the SalI restriction site located in the talin1 gene and the SacII restriction site present in the Clontech C1 vector.

### Antibodies and Reagents

Samples were fixed in 4% paraformaldehyde (PFA), warmed to 37°C, for 15 minutes before being washed thrice with PBS. For immunofluorescence, samples were permeabilized at room temperature with Triton X-100 (0.5%) for 5 minutes, before being washed thrice. The following primary antibodies were used at the indicated dilutions (in 1% BSA): anti-paxillin (clone 249, BD Transduction Labs, USA, 1:400), anti-vinculin (hVin1, Sigma, 1:400), 9EG7 (BD Biosciences, 1:200), anti-phosphotyrosine (4G10, Merck, 1:400). Actin was visualized using Texas-Red conjugated Phalloidin (Thermo Fisher), diluted 1:400. Secondary antibodies (Dylight 488-or 594-conjugated donkey anti-mouse or anti-rabbit) were purchased from Jackson ImmunoResearch and used at a dilution of 1:500. Y-27632 (Tocris Bioscience) was diluted in dH20 and used at a final concentration of 50 µM. Blebbistatin (Tocris Bioscience) and cytochalasin D (Sigma) were diluted in DMSO (Sigma) and used at a final concentration of 50 µM and 25 µg ml-1 respectively. Mitotracker Deep Red FM (Thermo Fisher) was dissolved in DMSO to a concentration of 1 mM. Prior to use, the stock was diluted in pre-warmed medium at a final concentration of 200 nM, before being added directly to cells 30 minutes prior to imaging. Mutagenesis was performed using the QuikChange Lightning site-directed mutagenesis kit (Agilent) according to the manufacturer’s instructions.

### Microscopy

Fluorescence loss after photobleaching (FLAP) experiments were performed as described previously (Stutchbury et al., 2017) using a spinning disk confocal microscope (CSU-X1, Tokogawa) supplied by Intelligent Imaging Innovations, Inc. (3i) equipped with a 63×/1.42 Plan Apo oil objective (Zeiss). One hour prior to imaging the medium was changed to pre-warmed Ham’s F-12 medium supplemented with 25 mM HEPES buffer, 1% FCS, 1% penicillin/streptomycin and 1% L-glutamine. PAGFP-tagged proteins were excited with a 405nm laser for 5 ms and post-activation images were acquired every 5 seconds for 5 minutes. The intensities of the post-activated PAGFP at mitochondria were measured manually using imageJ. Values were normalised to the intensity of the first post-activation image. Graphs were prepared using Prism 7.0 (Graphpad). Images of fixed samples were acquired using a Zeiss AxioObserver Z1 wide-field microscope equipped with a 100×/1.4 NA oil objective and an Axiocam MRm camera.

### Live-cell imaging

Images of TalinKO cells expressing GFP-Paxillin or GFP-TalinFL with RFP-LifeAct were acquired on a spinning disk confocal microscope (CSU-X1, Tokogawa) supplied by Intelligent Imaging Innovations, Inc. (3i) equipped with a heated stage maintained at 37°C, using a 100×/1.45 Plan Apo oil objective (Zeiss). One hour prior to imaging the medium was changed to pre-warmed Ham’s F-12 medium supplemented with 25 mM HEPES buffer, 1% FCS, 1% penicillin/streptomycin and 1% L-glutamine, with 5mM Mn^2+^ added as appropriate. Cell edge tracing was performed with the QuimP plugins for FIJI (Baniukiewicz et al., 2018), using the signal from the RFP-LifeAct channel.

### Adhesion curvature quantification

Images were acquired using an Olympus IX83 inverted microscope (Olympus) equipped with a 60×/1.42 PlanApo oil objective (Olympus), using green and red Lumencor LED excitation and the Sedat filter set (Chroma 89000). Images were collected using a Retiga R6 camera (Q-Imaging). Adhesion curvature was calculated by first generating a binary image of the adhesions, which was then skeletonized. The Analyze Skeleton plugin for FIJI was used to extract the number of pixels and Euclidean distance for each line. Adhesion curvature was quantified for lines above 0.2 µm by dividing the total length by the Euclidean distance.

### Traction Force microscopy

Traction forces were quantified by preparing polyacrylamide hydrogels containing 0.2 µm diameter red fluorescent beads (FluoSpheres carboxylate-modified red [580/605], Molecular Probes) as described previously (Atherton et al., 2015). TalinKO cells expressing either GFP-TalinFL or GFP-Paxillin were allowed to spread on the hydrogels in the presence of 5mM Mn^2+^ for 1 hour. Images were acquired of the cells and the beads under strain using an Olympus IX83 inverted microscope with a heated stage maintained at 37°C, with a 100× UPlanFL 100×/0.17 objective, using green and red Lumencor LED excitation and a Sedat filter set (Chrome [89000]). Images were collected using a Retiga R6 camera (Q-Imaging). Cells were detached by adding 1% Triton X-100 for 45 minutes, before images of the beads without strain were acquired. After aligning the stressed and relaxed bead images to correct for drift, the deformation of the hydrogel was calculated using PIV plugins for ImageJ (Tseng et al., 2012). The total force was calculated by measuring the integrated density of the magnitude maps, using the whole cell area as a mask.

---

**Fig. S1.**
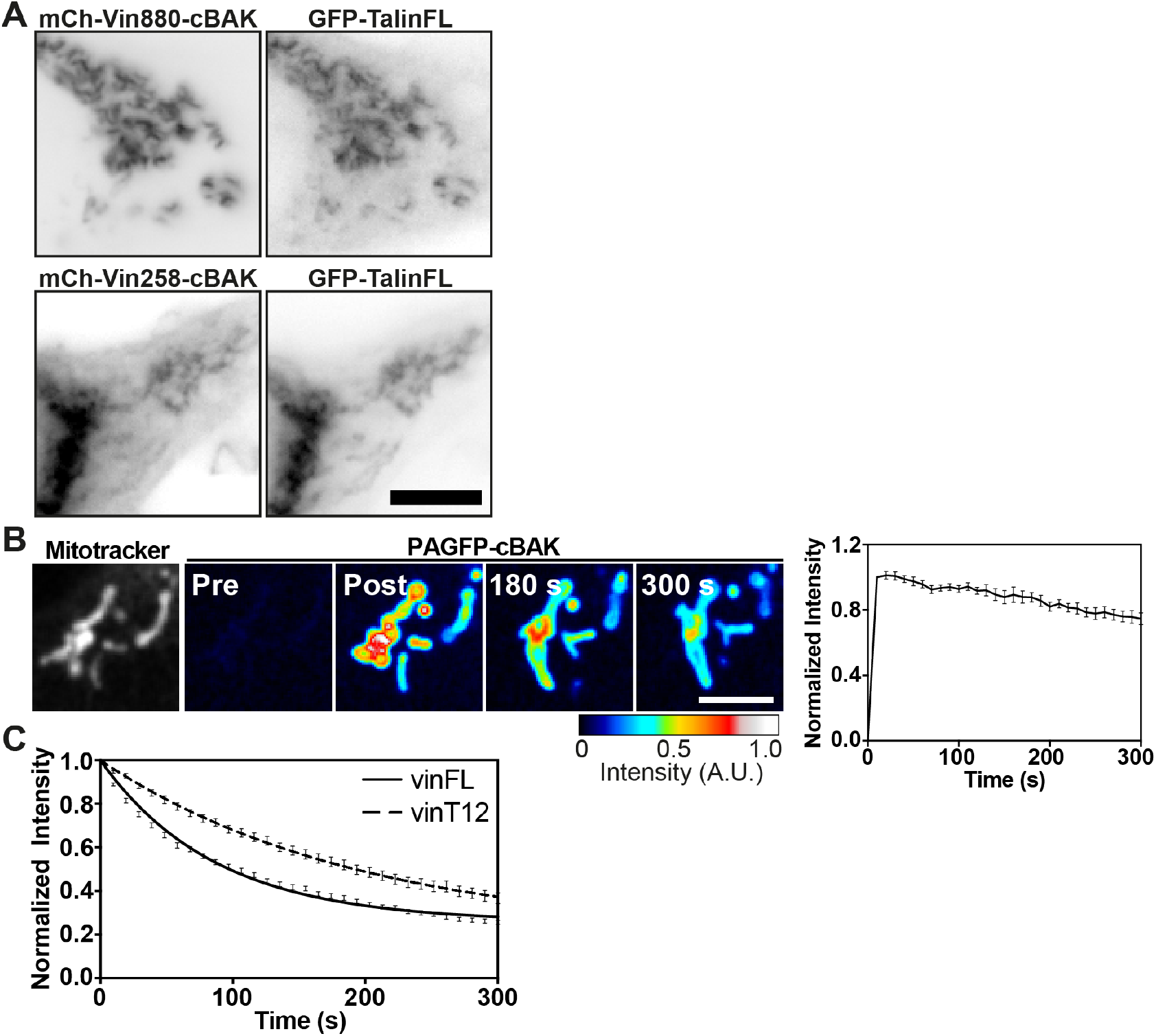
Recruitment of Talin to truncated vinculin constructs. **A**. Co-expression of GFP-TalinFL with either mCh-vin880-cBAK (lacking the C-terminal vinculin tail) or mCh-vin258-cBAK (the D1 domain of vinculin only) in NIH3T3 cells shows that both cBAK constructs can recruit talin. Scale bar indicates 10 µm. **B**. FLAP curve and images of PAGFP-cBAK at mitochondria marked using MitoTracker Deep Red FM in NIH3T3 cells reveals this construct stably integrates into the mitochondrial membrane. Scale bar indicates 5 µm, error bars are S.E.M, n = 15 mitochondria from 5 cells; results are representative of 3 independent repeats. **C**. FLAP curves of PAGFP-TalinFL at FAs co-expressed with either mCh-vinFL or mCh-vinT12. Note the reduced turnover of talin at FAs when co-expressed with vinT12 Error bars are S.E.M, n = 92 (vinFL) or 68 (vinT12) FAs, from 10 – 15 cells. Data are pooled from 3 independent experiments.

**Fig. S2.**
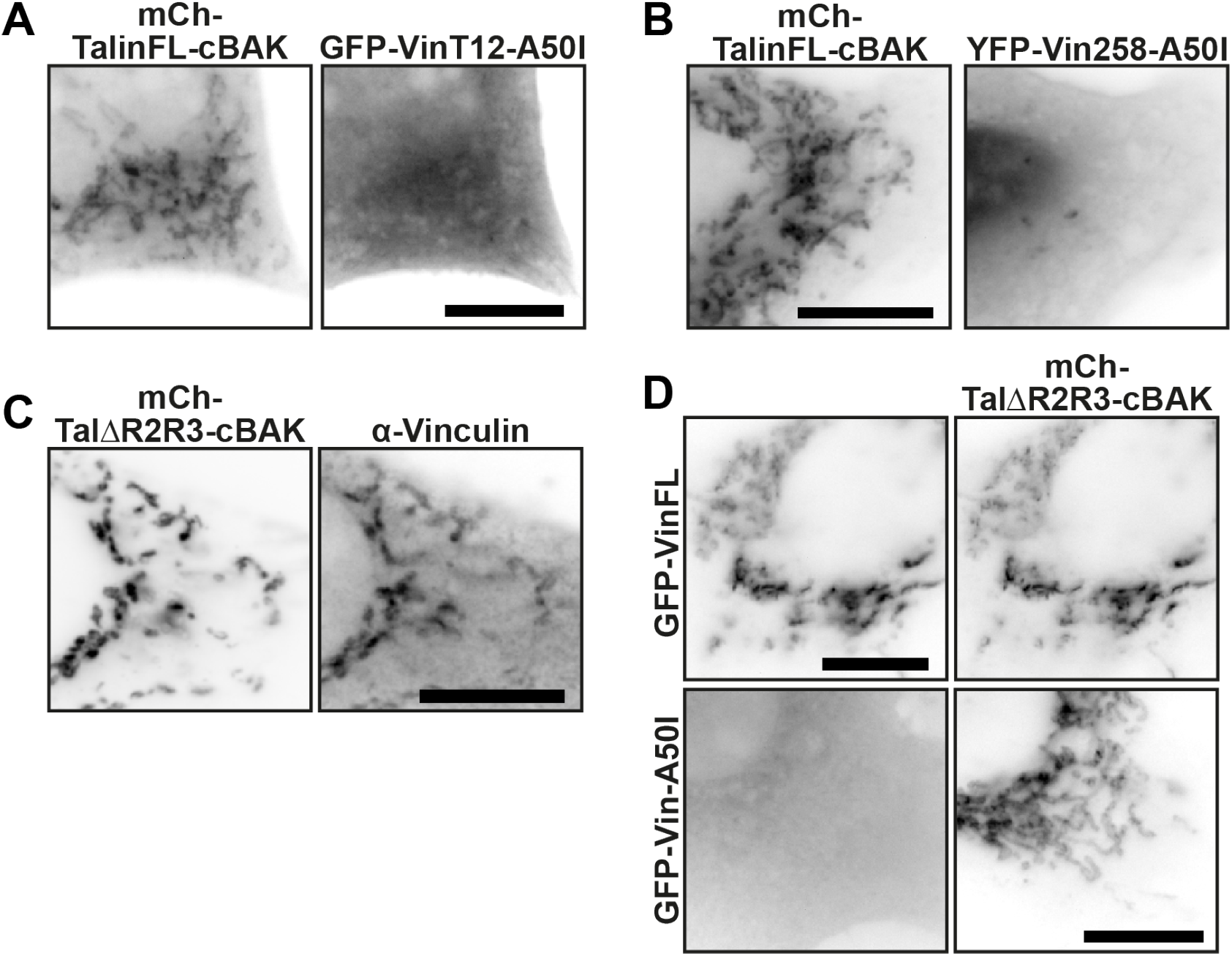
Force-independent interactions between active vinculin and talin, and vinculin and active talin are direct. **A, B**. mCh-TalinFL-cBAK is unable to recruit activated forms of vinculin containing an A50I mutation in the canonical talin binding site in the vinculin head domain (D1)) (A) GFP-vinT12-A50I, or (B) YFP-Vin258-A50I. **C**. An activated talin mutant targeted to mitochondria (mCh-Tal∆R2R3-cBAK) recruits endogenous vinculin (hVin1 antibody staining) in NIH3T3 cells. **D**. Mutating the canonical talin binding site within the D1 domain of full-length vinculin (GFP-Vin-A50I) blocks the recruitment of vinculin to active talin at mitochondria (mCh-Tal∆R2R3-cBAK). Scale bars in A, B, C, D indicate 10 µm.

**Fig. S3.**
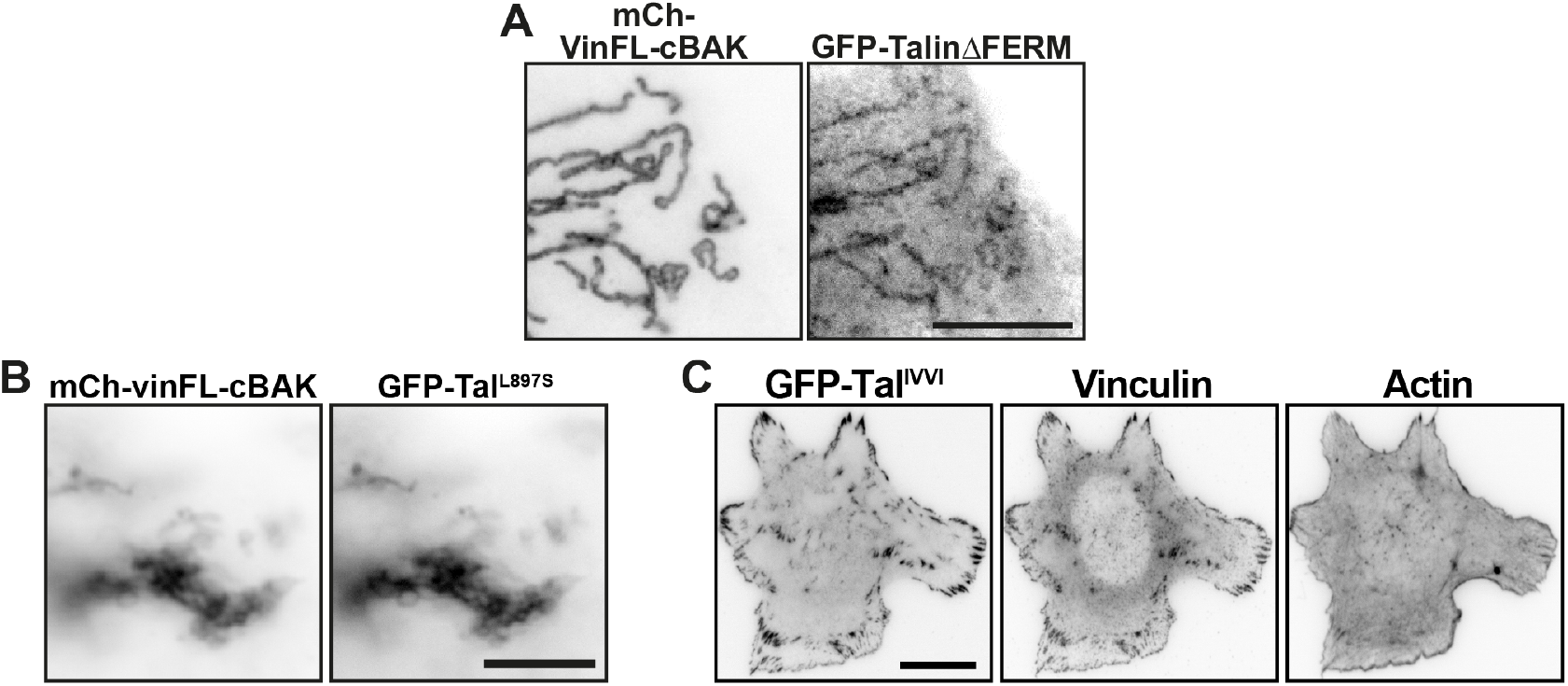
Localisation of Talin^IVVI^ to FAs in TalinKO cells. **A**. An activated GFP-Talin∆FERM construct is recruited to mitochondrial-targeted mCh-vinFL-cBAK in NIH3T3 cells. Scale bar indicates 10 µm. **B**. GFP-Tal^L897S^ containing a destabilising mutation in the R3 domain [31] is recruited to mCh-vinFL-cBAK when co-expressed in NIH3T3 cells; scale bar indicates 10 µm. **C**. GFP-Tal^IVVI^ containing an R3 stabilising mutation localises to FAs when expressed in Talin1-/2-(TalinKO) cells.

**Fig. S4.**
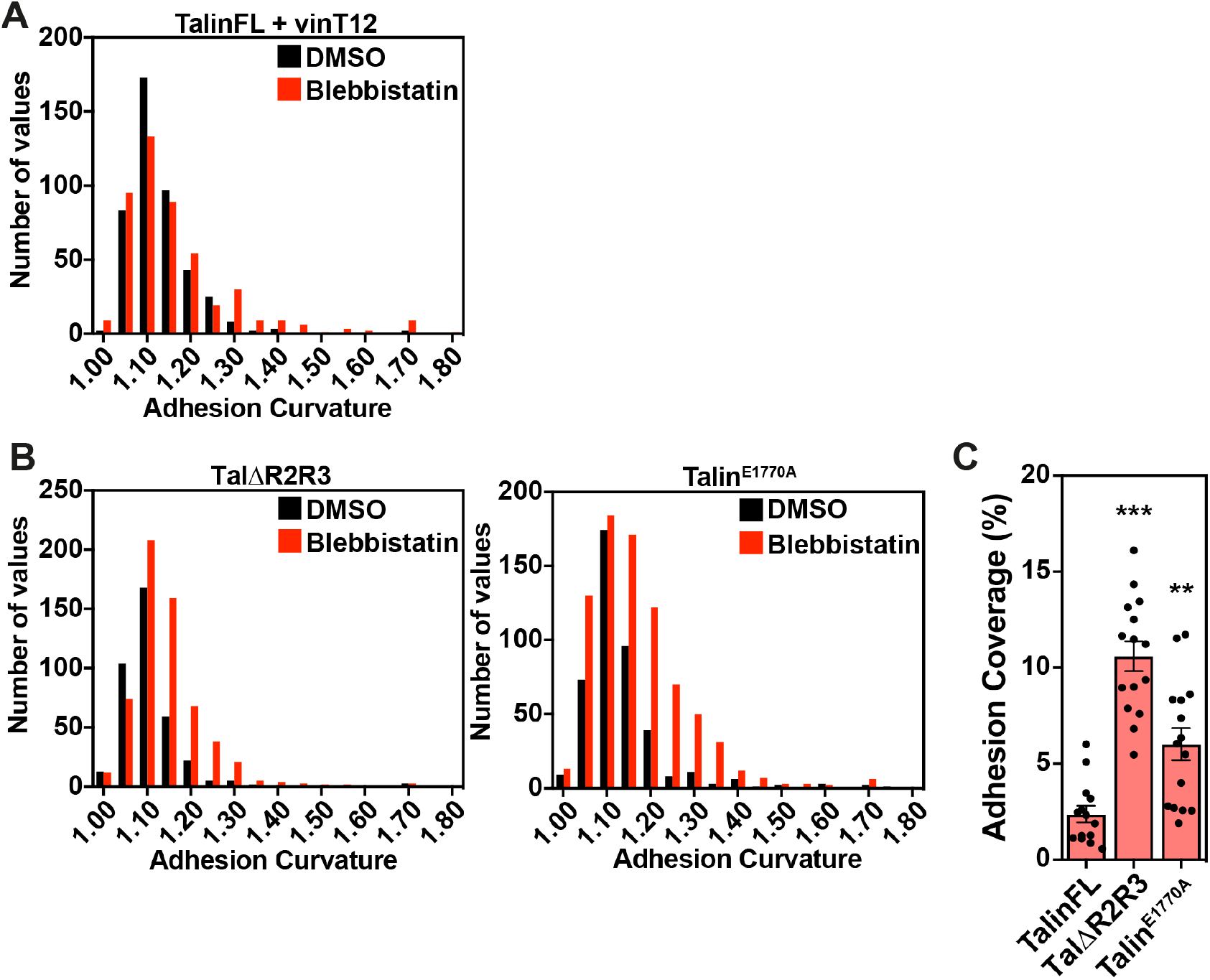
Active talin constructs form disorganised adhesions when spreading in the absence of intracellular tension. **A**. Histograms of adhesion curvature calculated from talinKO cells co-expressing GFP-TalinFL and mCh-vinT12 treated in suspension with either DMSO or blebbistatin. **B**. Histograms of adhesion curvature calculated from talinKO cells expressing GFP-Tal∆R2R3 or GFP-Talin^E1770A^, after DMSO or blebbistatin treatment. **C**. Quantification of the percentage of the cell consisting of adhesions (quantified from the GFP signal) in TalinKO cells expressing GFP-Tal∆R2R3 or GFP-Talin^E1770A^. Cells were pre-treated in suspension with blebbistatin (50 µM) or an equivalent volume of DMSO, for 45 minutes, before being seeded onto fibronectin-coated glass and fixed after 15 minutes of spreading. Graphs show the mean and S.E.M; results are representative of 3 independent experiments, ** indicates p<0.01; *** indicates p<0.001, significance against TalinFL.

**Fig. S5.**
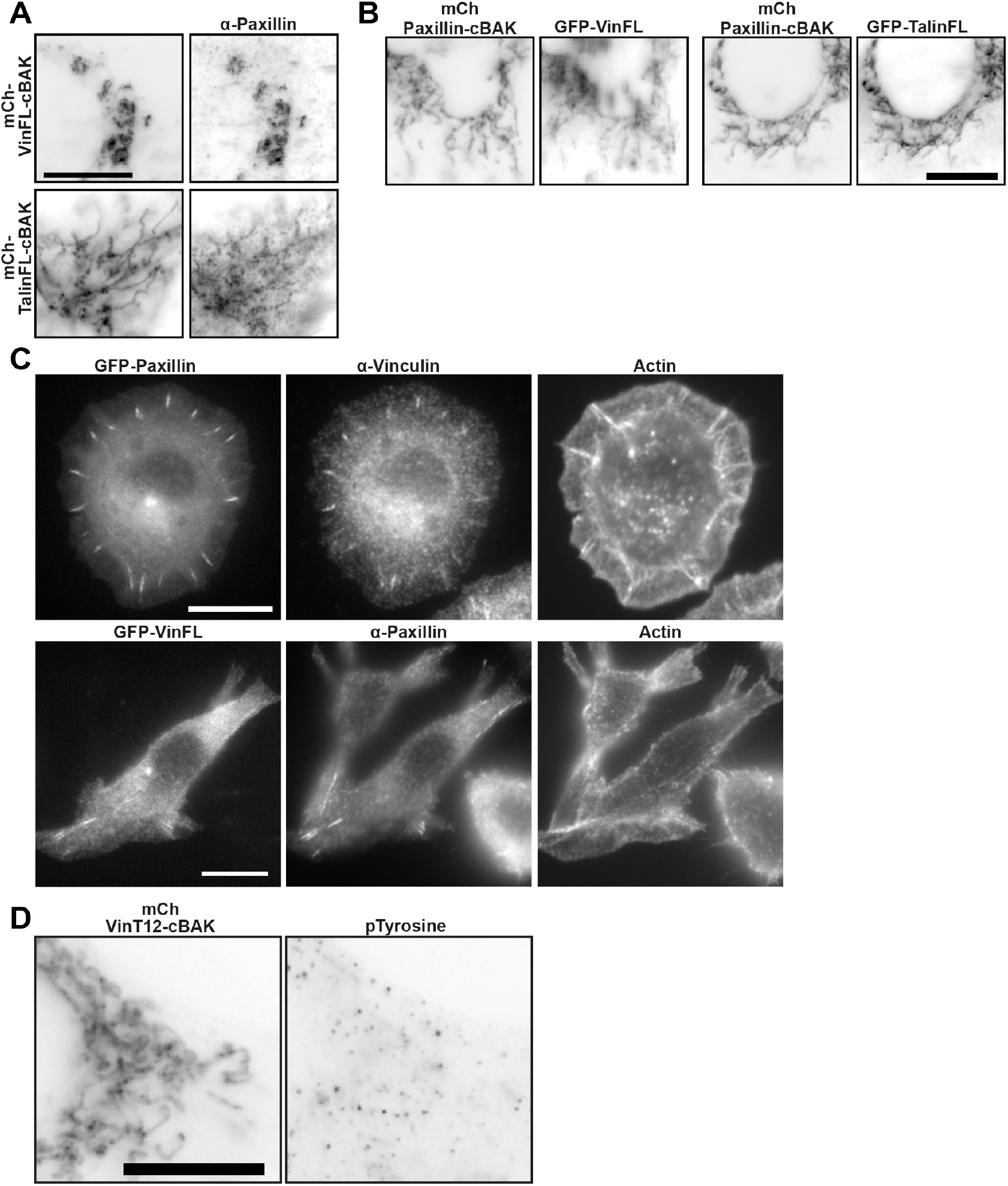
Paxillin can recruit inactive vinculin and talin independent of forces. **A**. Endogenous paxillin is recruited to both inactive vinFL-cBAK inactive talinFL-cBAK. **B**. mCh-Paxillin-cBAK is able to recruit either GFP-vinFL of GFP-talinFL to mitochondria. **C**. TalinKO cells expressing either GFP-Paxillin or GFP-VinFL were spread on fibronectin for one hour in the presence of Mn^2+^ (5 mM) before fixation and staining for endogenous vinculin or paxillin, respectively. **D**. Representative image of an NIH3T3 cell expressing mCh-vinT12-cBAK stained for phosphotyrosine. Scale bars in A-D indicate 10 µm.

